# DRB1 as a mediator between transcription and microRNA processing

**DOI:** 10.1101/2019.12.30.890665

**Authors:** Dawid Bielewicz, Jakub Dolata, Mateusz Bajczyk, Lukasz Szewc, Tomasz Gulanicz, Susheel Sagar Bhat, Anna Karlik, Artur Jarmolowski, Zofia Szweykowska-Kulinska

## Abstract

DRB1 (HYL1) is a double-stranded RNA binding protein involved in miRNA processing in plants. It is a core component of the Microprocessor complex and enhances the efficiency and precision of miRNA processing by DCL1 protein. In this work, we report a novel function of DRB1 protein in the transcription of *MIR* genes. DRB1 co-localizes with RNA Polymerase II and affects its distribution along *MIR* genes. Moreover, proteomic experiments revealed that DRB1 protein interacts with many transcription factors. Finally, we show that the action of DRB1 is not limited to *MIR* genes as it impacts expression of many other genes, majority of which are involved in plant response to light. These discoveries add DRB1 as another player of gene regulation at transcriptional level, independent of its role in miRNA biogenesis.

## Introduction

MicroRNAs (miRNAs) are small ~21nt RNA molecules that play vital role in post-transcriptional regulation of gene expression (Bartel 2004). miRNAs are involved in a vast range of metabolic processes and hence act as regulatory molecules in overall plant development, for example in processes like determination of leaf shape, flowering time and response to environmental conditions (Palatnik, Allen et al. 2003, Chen 2004, Kruszka, Pieczynski et al. 2012, Barciszewska-Pacak, Milanowska et al. 2015). miRNA genes (*MIR*) are transcribed by RNA Pol II (Xie, Allen et al. 2005) and the primary transcripts (known as primary-miRNAs; pri-miRNAs) contain a stem-loop region. The pri-miRNAs are then cleaved to release mature miRNAs by a complex of enzymes known as the Microprocessor. Notably, the stem-loop region of pri-miRNAs serves as a mark for the Microprocessor to recognize and start processing the pri-miRNAs (Kurihara and Watanabe 2004, Dolata, Taube et al. 2018). Microprocessor is a complex of three core proteins, Dicer Like 1 (DCL1), Double-stranded RNA Binding protein 1 (DRB1) (also known as Hyponastic Leaves 1 (HYL1)) and Serrate (SE) (Yang, Liu et al. 2006, Fang and Spector 2007, Dong, Han et al. 2008, Dolata, Taube et al. 2018). DCL1 is a RNAse III type enzyme that cuts and releases miRNA/miRNA* from pri-miRNAs in a two-step reaction (Park, Li et al. 2002, Reinhart, Weinstein et al. 2002). DRB1 and SE assist and increase the accuracy of pri-miRNA cleavage by DCL1 (Kurihara, Takashi et al. 2006). All three of these core components are necessary for proper functioning of Microprocessor and in mutants of any of these proteins, pri-miRNAs accumulate and mature miRNAs are downregulated (Laubinger, Sachsenberg et al. 2008, Szarzynska, Sobkowiak et al. 2009, Zielezinski, Dolata et al. 2015). Two modes of Microprocessor mediated pri-miRNA cleavage have been described: base to loop (first cut at the base, second closer to the loop structure) and loop to base (first cut in the loop and second closer to the base), depending on the directionality of DCL1 cleavage (Kurihara and Watanabe 2004, Bologna, Mateos et al. 2009).

DRB1, a double stranded RNA binding protein, contains two double stranded RNA binding domains (dsRBDs) in its N-terminal region followed by a nuclear localization signal (NLS) and an unstructured C-terminal region (Yang, Chen et al. 2010). DRB1 is considered to form a homodimer and bind the miRNA/miRNA* duplex region of pri-miRNA (Burdisso, Milia et al. 2014, Yang, Ren et al. 2014). DRB1 knockout has a pleiotropic impact on the phenotype of plants (among the others defects in leaf shape and late flowering time) and their response to auxin and ABA (Lu and Fedoroff 2000, Vazquez, Gasciolli et al. 2004). So far, no ortholog of DRB1 has been identified in mammalian cells. However, similar to plants, mammalian RNA nucleases (Drosha and Dicer) also require proteins containing dsRBDs: DiGeorge syndrome critical region 8 (DGCR8) and transactivation response element RNA-binding protein (TRPB), for proper functioning (Han, Lee et al. 2004, Chendrimada, Gregory et al. 2005, Haase, Jaskiewicz et al. 2005). Both, DRB1 and DGCR8 are nuclear proteins involved in first step of pri-miRNA maturation i.e. cleavage by RNAse III (Gregory, Yan et al. 2004). However, like TRBP, DRB1 can be phosphorylated which affects its activity (Paroo, Ye et al. 2009, Manavella, Hagmann et al. 2012, Achkar, Cho et al. 2018).

miRNA production in mammalian cells is co-transcriptional as Drosha and DGCR8 have been shown to bind regions in the proximity of many gene promoters (not just *MIR* genes) and consequently Drosha knockdown in HeLa cells negatively impacts gene transcription (Gromak, Dienstbier et al. 2013). Similarly, in plants, co-transcriptional processing of plant pri-miRNAs has been proposed through the association of DCL1 with the chromatin and Elongator complex (Fang, Cui et al. 2015). However, presence of pri-miRNA transcripts is necessary for the association of DCL1 with chromatin. Since the interaction between DCL1 and DRB1 has been established, here we ask whether it possible that DRB1 is also involved in the transcription of *MIR* genes?

## Results

### DRB1 is a positive regulator of *MIR* gene transcription

To investigate the possibility that DRB1 protein is involved in the transcription of *MIR* genes we used a GUS reporter line system. We independently crossed two reporter lines with GUS under two different *MIR* gene promoters (p*MIR393A*:GUS and p*MIR393B*:GUS (Parry, Calderon-Villalobos et al. 2009)) in the wild type (Col-0) background with *hyl1-2* mutants (DRB1 knockout mutants). We observed that the expression of GUS protein driven by *MIR* gene promoter was remarkably lower in *hyl1-2* background as compared to Col-0 (FIG. 1A). We measured the level of GUS transcripts in p*MIR393A*:GUS, p*MIR393B*:GUS, p*MIR*392A:GUSx*hyl1-2*, and p*MIR*392B:GUSx*hyl1-2* reporter lines using RT-qPCR and observed a significant decrease in the level of GUS transcripts in p*MIR*392A:GUSx*hyl1-2*, and p*MIR*392B:GUSx*hyl1-2* plants. Additionally, we also tested the levels of pri-miRNA393A and pri-miRNA393B in the obtained lines and as expected, these pri-miRNAs were upregulated in the *hyl1-2* mutant background (Fig. 1B). These data indicate that DRB1 which is involved in pri-miRNA processing may also regulate the transcription of *MIR* genes. However, it is also possible that the strong downregulation of GUS expression in the *hyl1-2* background is caused by a feedback mechanism in which the global downregulation of miRNAs in *hyl1-2* results in an upregulation of a yet unidentified transcriptional factor(s) that in turn inhibits *MIR393A* and *MIR393B* transcription. To address this question, we decided to restore the miRNA levels in the reporter lines within *hyl1-2* background. For this purpose we crossed these lines with a *DCL1* mutant: *dcl1-13* (Tagami, Motose et al. 2009). *dcl1-13* mutant has a point mutation in the *DCL1* gene which promotes DCL1 activity in absence of DRB1. The point mutation in *DCL1* gene results in an amino acid substitution of Glu to Lys in the ATPase/DExH-box RNA helicase domain. Hence, the expression of *dcl1-13* allele in the *hyl1-2* background restores the level of miRNAs and rescues plants from the developmental abnormalities associated with lower level of miRNAs. We prepared a *hyl1-2/dcl1-13* transgenic line carrying genomic sequence of mutated *DCL1* gene under its native promoter. We obtained four independent transgenic lines and confirmed previous results showing that the expression of *dcl1-13* allele restores the levels of pri-miRNAs and mature miRNAs in *hyl1-2* background (Fig. S1). We then crossed our reporter lines with *hyl1-2*/*dcl1-13* transgenic plants. Analysis of GUS staining in the offspring clearly showed that GUS protein is still inefficiently expressed in the *hyl1-2*/*dcl1-13* mutant background (Fig. 1A, right panel). Additionally, we also performed RT-qPCR analysis to measure the GUS transcript level and like in *hyl1-2* mutant, in *hyl1-2*/*dcl1-13* mutant plants, GUS transcript was downregulated in comparison to wild type plants (FIG. 1B, right panel). Hence, our data shows that DRB1 is a direct positive regulator of *MIR* gene transcription.

**Figure 1.**
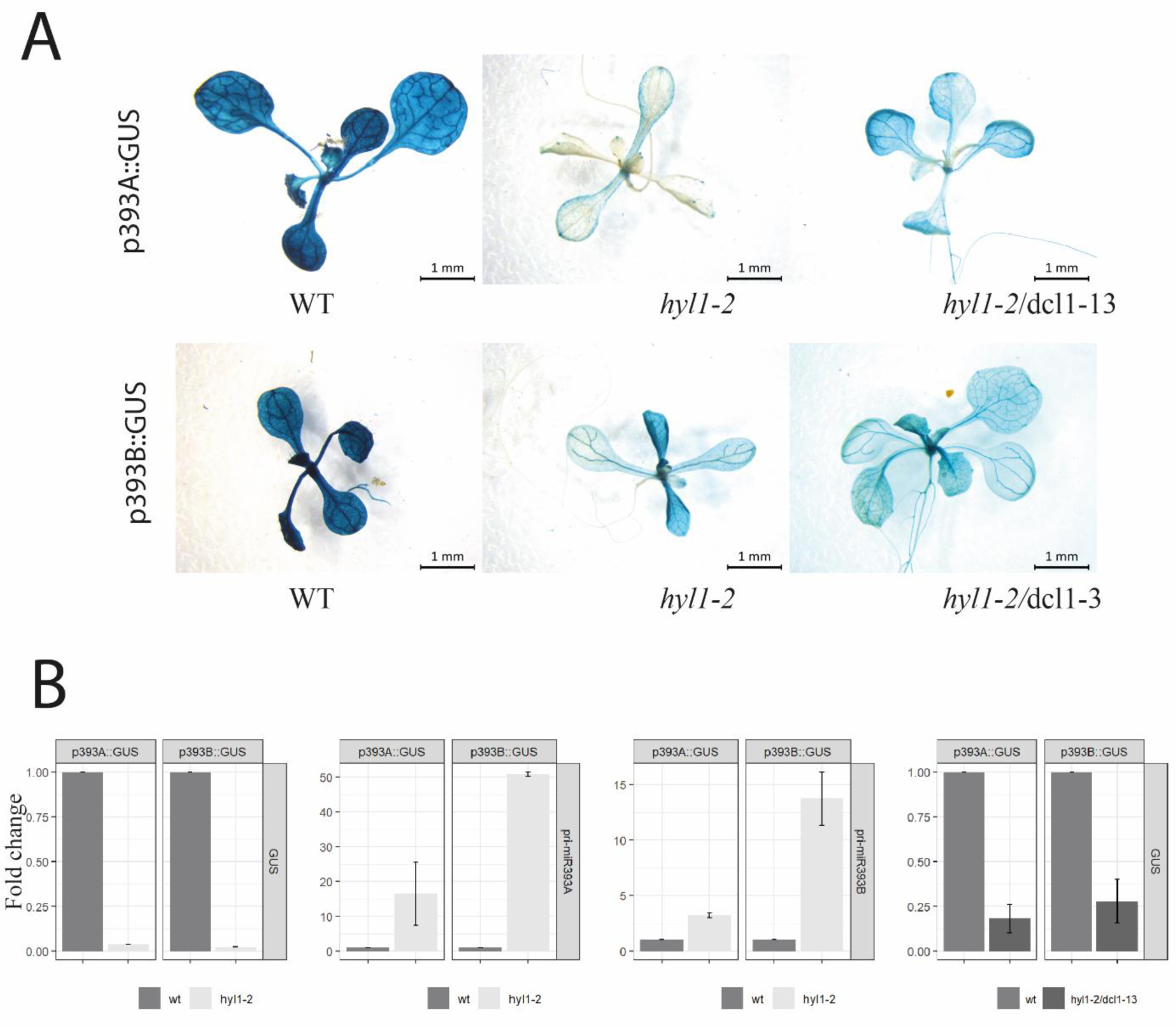
DRB1 is a positive factor for transcription of *MIR* genes. **a)** Images show GUS staining of seedlings reporter lines in wild type (left), *hyl1-2* (middle) and *hyl1-2/dcl1-13* (right) backgrounds. **b)** RT-qPCR analysis of GUS transcript and pri-miRNA levels in the reporter lines in wild type and *hyl1-2* and *hyl1-2/dcl1-13* backgrounds. Error bars indicate SD (n=3).

### DRB1 colocalizes with RNA polymerase II

The observation that DRB1 is required for a proper transcription of *MIR393A* and *MIR393B* prompted us to test the stage at which it affects transcription. We tested the co-localization of DRB1 with RNA Pol II at different stages of transcription. We performed immunolocalization in fixed nuclei from wild type plants and observed strong colocalization of DRB1 and total RNA Pol II (Fig. 2). We used antibodies specific for Serine5 (transcription initiation) and Serine2 (transcription elongation) of RNA Pol II C-terminal domain (CTD) independently to further characterize the nature of these interactions. The results showed that DRB1 is associated with RNA Pol II already at the transcription initiation stage (Ser5) and remains associated with RNA Pol II during the elongation step (Ser2). Since DRB1 is a part of Microprocessor core complex we considered the possibility that DRB1 is brought to RNA Pol II by another member of Microprocessor. Recently it was shown that SE is directly associated with specific regions of Arabidopsis chromatin (Speth, Szabo et al. 2018). Apart from its role in miRNA biogenesis, SE is involved in many processes connected with RNA metabolism like splicing, 3’-end formation, RNA transport and RNA stability (Laubinger, Sachsenberg et al. 2008, Raczynska, Stepien et al. 2014). To exclude the possibility of SE mediating DRB1 and RNA Pol II co-localization, we investigated the interaction of DRB1 with RNA Pol II in the *se-2* mutant plants. (Fig S2). Similar to our previous experiment, these results were also obtained for total RNA Pol II and for RNA Pol II phosphorylated at Ser5 or Ser2. We did not see any decrease in co-localization of DRB1 and RNA Pol II in the absence of SE, thus showing that the association of DRB1 with RNA pol II is not SE dependent. We also addressed the question whether DRB1 is involved throughout the whole process of transcription. For this purpose, we performed immunolocalization studies for DRB1 and RNA Pol II using antibodies against DRB1 and antibodies which recognize RNA Pol II phosphorylated at threonine 4 (Thr4) of its CTD domain (mostly associated with transcription termination or 3’ end processing events) (Hintermair, Heidemann et al. 2012, Nojima, Rebelo et al. 2018). Our co-localization results showed that DRB1 did not co-localize with RNA pol II phosphorylated at Thr4 in either wild-type or *se-2* mutant plants. These data show that DRB1 is involved in transcription process from initiation to elongation, but not at the termination stage and DRB1’s role in transcription stimulation is not mediated by other proteins, at least not by SE.

**Figure 2.**
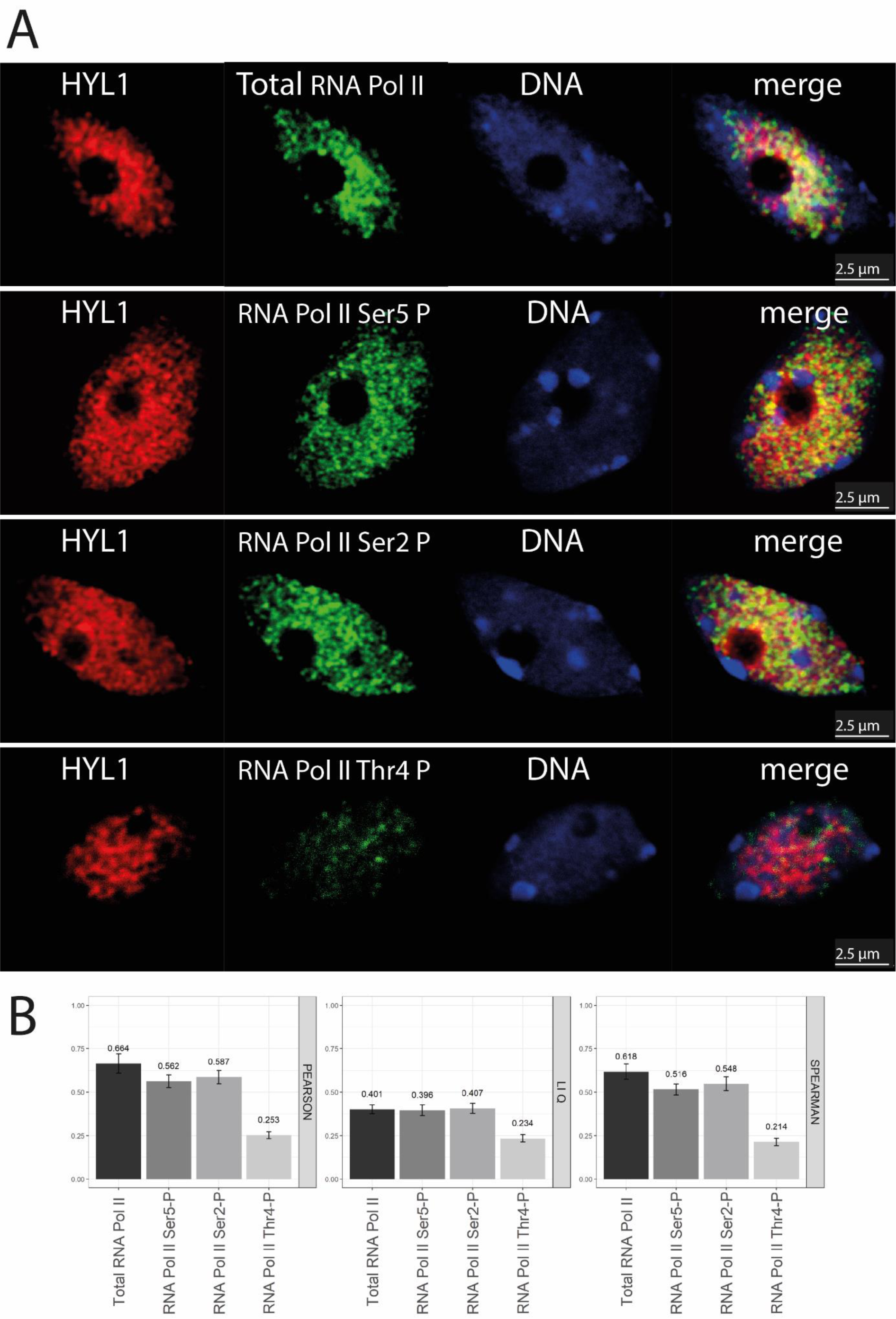
DRB1 colocalizes with RNA polymerase II in wild type plants: **a)** Nuclei from fixed cells where DRB1 is shown in red, RNA Pol II in green and DNA in blue. The merge column shows all 3 channels. **b)** Colocalization scores of DRB1 and RNA Pol II calculated by three different approaches (Pearson, Spearman and LiQ).

### DRB1 protein interacts with transcription factors

Since we found that DRB1 acts as a positive regulator of *MIR* gene transcription and co-localizes with RNA Pol II we decided to test whether it also interacts with other known transcription factors. For this experiment, we used a transgenic line containing DRB1 tagged with HA epitope (*pDRB1:DRB1:HA*) and anti-HA antibodies to co-immunoprecipitate with its potential protein interactors which were later identified by Mass Spectrometry analysis. As a benchmark for our experiment, we searched for previously published data regarding DRB1 interactors and compared with our data. We were able to identify bona fide DRB1 partners KETCH1 and CDC5 in our data (Fig 3). (Zhang, Xie et al. 2013, Zhang, Guo et al. 2017). KETCH1 is a karyopherin enabling the transport of DRB1 to nucleus and CDC5 is a MYB-transcription factor. Interestingly CDC5 was showed to be involved in the transcription of *MIR* genes (Zhang, Xie et al. 2013). Similar to DRB1 mutant plants, in the CDC5 knockout mutant the GUS reporter which was under the control of *MIR* gene promoter was downregulated. However, *cdc5* mutants showed no enrichment in pri-miRNA level, which can be explained that CDC5 is involved in transcription of *MIR* genes but not in the processing of pri-miRNA like DRB1 (Vazquez, Gasciolli et al. 2004, Zhang, Xie et al. 2013). Interestingly, in our co-IP results we found that DRB1 was associated also with transcription factors from the Topless transcription factor family (Fig 3). However, the interaction between Topless transcription factors and DRB1 requires more detailed investigation.

**Figure 3.**
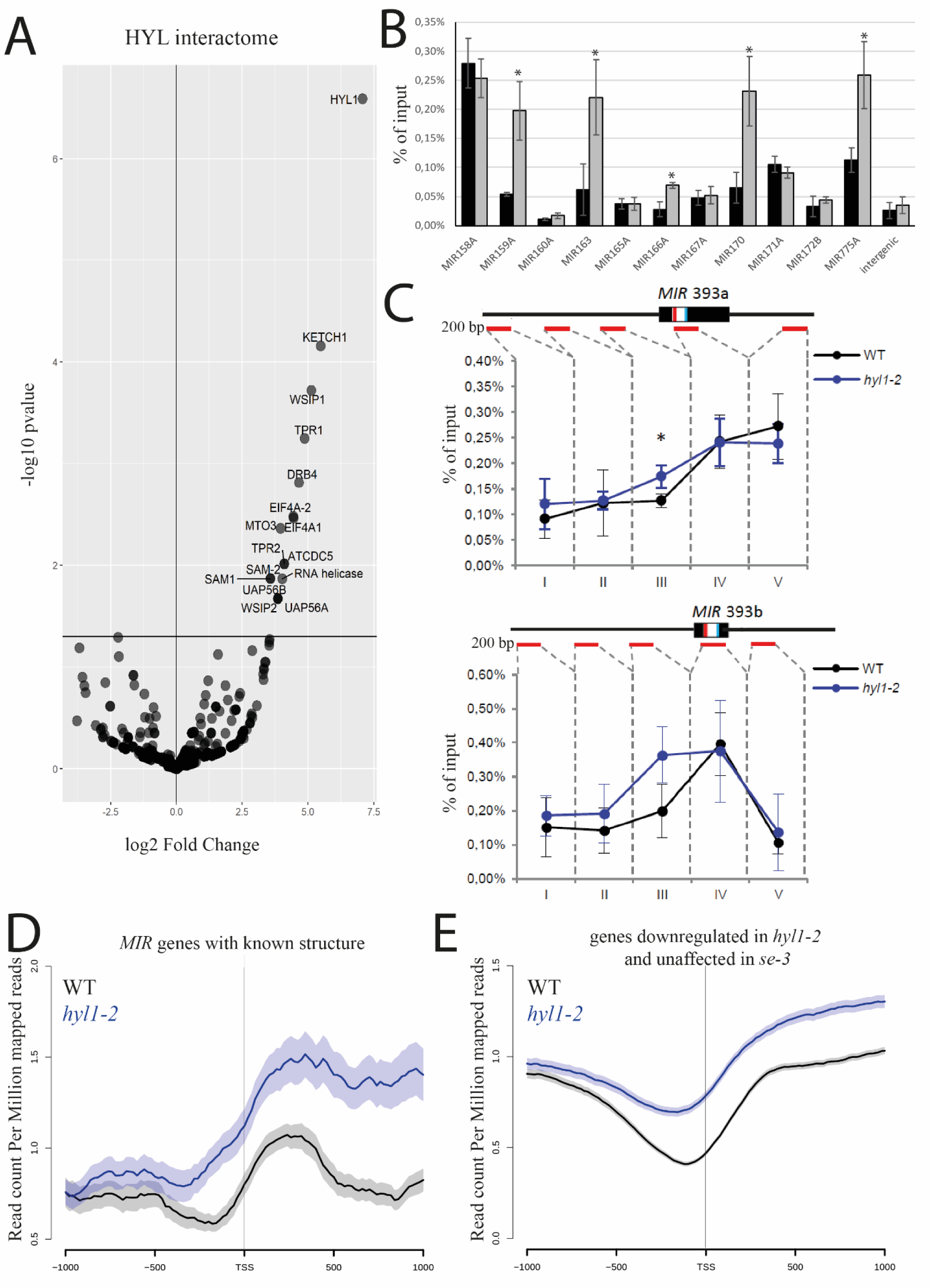
DRB1 interacts with transcriptional factors and is important for proper distribution of RNA polymerase II along *MIR* genes: **a)** The DRB1 interactome determined by co-immunoprecipitation followed by Mass Spectrometry **b)** The occupancy of total RNA Pol II at promoter regions on different *MIR* genes (~200 bp upstream of TSS). **c)** The occupancy of total RNA Pol II at the *MIR393A* or *MIR393B* loci using ChIP followed by qPCR. The region marked with an asterisk represents the statistically significant enrichment of RNA Pol II in *hyl1-2* as compared to wild-type plants. Schematic gene structure is shown, and red lines show the amplified regions. Error bars represent the SD of three independent biological replicates.

### Distribution of RNA polymerase II is affected in *hyl1-2* mutant

In light of our data showing that DRB1 and RNA Pol II co-localize, we tested whether this co-localization also affects RNA Pol II occupancy on *MIR* genes. For this purpose, we performed Chromatin Immunoprecipitation (ChIP) using antibody against total RNA Pol II followed by qPCR in wild-type and *hyl1-2* mutants. We compared the occupancy of RNA Pol II on selected *MIR* genes (~200 bp upstream of transcription initiation start site) in *hyl1-2* and wild-type plants (Fig. 3B). We selected the *MIR* genes which were tested previously in the *cdc5* mutant plants (Zhang, Xie et al. 2013). The results showed that RNA Pol II distribution is affected at several *MIR* genes promoter regions. Our data showed an increased accumulation of RNA Pol II at the region approximately 200 base pair upstream from the transcription initiation site (TSS) for few tested *MIR* genes in the *hyl1-2* mutant when compared to wild type plants. This observation is opposite to what was reported in *cdc5* mutant plants. Additionally, we examined the RNA Pol II distribution in detail on *MIR393A* and *MIR393B* genes (Fig. 3C). The higher occupancy of RNA polymerase II in these regions may suggest that, in the *hyl1-2* mutant: 1) transcription of selected *MIR* genes is more efficient; or 2) transition from initiation to transcription elongation is deregulated. Keeping in mind our previous results where transcription of GUS reporter was downregulated in a *hyl1-2* mutant when compared to wild-type plants we suggest that the accumulation of RNA Pol II on the promoter region of *MIR* genes in *hyl1-2* is a result of RNA Pol II’s inability to transition from initiation to elongation. Furthermore, we performed ChIP-seq experiments using antibodies against total RNA pol II in wild-type and *hyl1-2* mutant plants for a global analysis of RNA pol II distribution along *MIR* genes. For the global ChIP-seq analysis we selected only *MIR* genes which are independent transcriptional units with determined TSS (Bielewicz, Dolata et al. 2012, Zielezinski, Dolata et al. 2015). Results clearly show increased RNA Pol II occupancy in the region of TSS as well as in the gene body of *MIR* genes in *hyl1-2* mutant in comparison to wild-type plants (Fig. 3D). ChIP-seq data also confirmed the results obtained for *MIR*393A gene (Fig. S3A). Our data suggests that DRB1 acts a positive factor for RNA Pol II to transition through initiation to elongation.

### Knockdown of DRB1 affects gene expression of many genes

We were interested whether DRB1 could also affect transcription of other genes transcribed by RNA Pol II. To investigate this possibility, we analyzed publicly available transcriptomic data obtained from wild-type and *hyl1-2* mutant plants. Theoretically, in *hyl1-2* mutant, a generally low level of miRNAs should be accompanied by a general upregulation of miRNA targets. On the contrary, we noticed that approximately half (1531 genes) of differentially expressed genes (DEGs) (3036 genes) in *hyl1-2* mutants are downregulated as compared to wild-type plants (Fig. 4A). We then compared these results with the transcriptomic data obtained from *se-3* mutant. Our analysis showed that almost half of the DEGs are common between *se-3* mutant and *hyl1-2* mutant, indicating that the functions of DRB1 and SE proteins overlap (Fig. 4B). Interestingly, there is a big group of DEGs which are specifically downregulated in *hyl1-2* mutant plants (769 genes). In agreement with our ChIP-seq data, RNA Pol II occupancy (on TSS and gene body) was markedly increased in genes (not limited to *MIR* genes) whose transcription was affected only by DRB1 (Fig. 3E). Gene ontology analysis showed that proteins encoded from these genes localized mostly in chloroplasts (Fig. 4C). Additionally, the biggest group of genes downregulated in *hyl1-2* mutant is involved in biological processes of ‘plastid organization’ according to our gene ontology analysis (Fig. S3B). It was known that light is an important factor for DRB1 maintenance in the cell (Cho, Ben Chaabane et al. 2014). In the night DRB1 is degraded by yet unidentified protease which results in downregulation of mature microRNA level. However, our analysis suggests that DRB1 may be involved in plant response to light independent of its role in the miRNA pathway. Our data regarding the role of DRB1 outside of miRNA biogenesis is further supported by a recently published work that uncovered a novel function of DRB1 connected to skotomorphogenesis independent of its role in miRNA biogenesis (Sacnun, Crespo et al. 2019). Taken together, our data point towards a role of DRB1, outside of miRNA biogenesis, where it acts as a general transcriptional factor, though a lot more work needs to be done before we can completely elucidate the mechanism of transcriptional gene regulation by DRB1.

**Figure 4.**
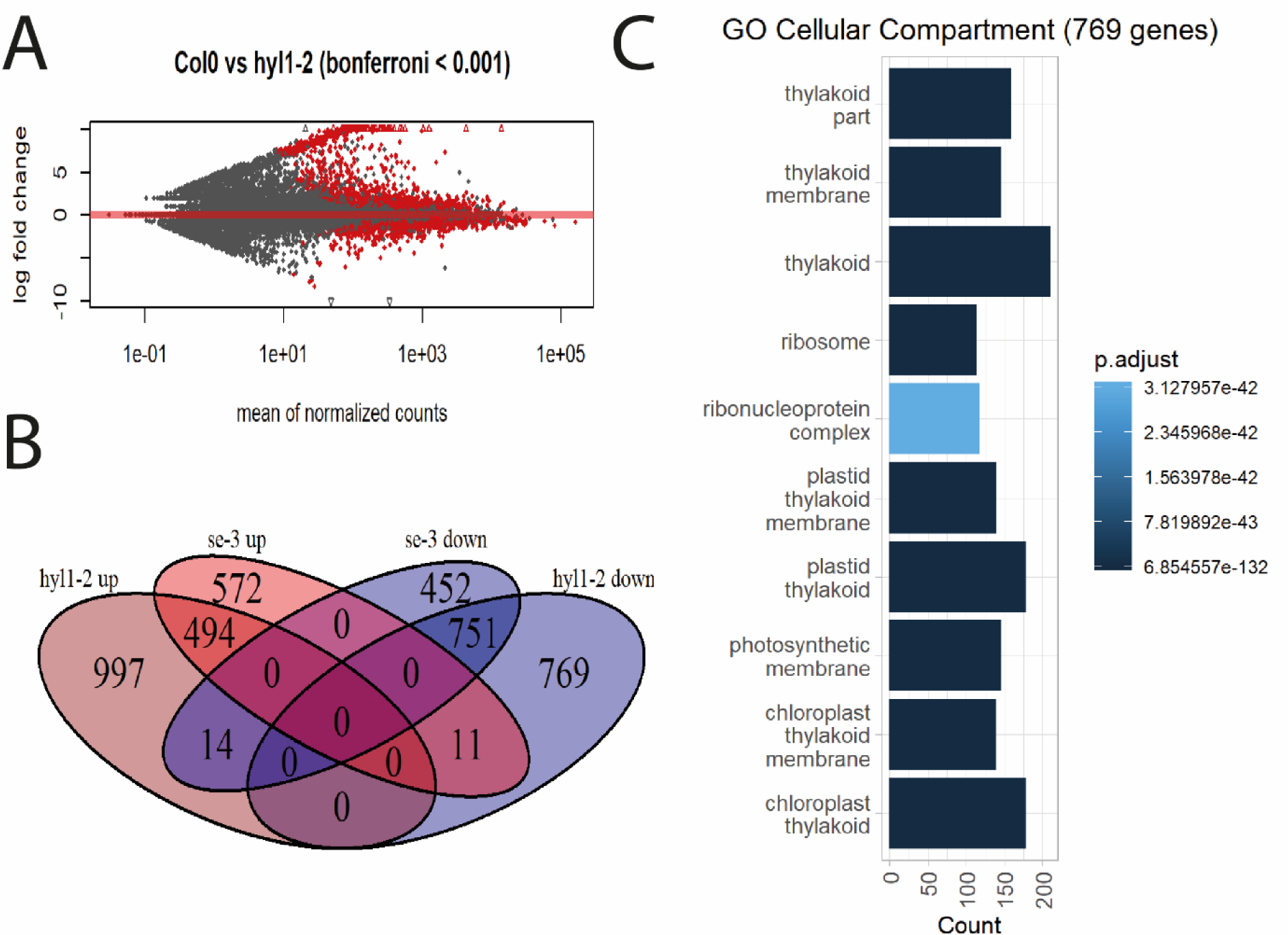
Downregulation of RNA Pol II transcripts in DRB1 mutant. **a)** MAplot showing the DEGs in *hyl1-2* mutant as compared to wild type. Red color indicates the statistically significant genes **b)** Venn diagram showing the overlap of DEGs between *hyl1-2* and *se-3* mutants. **c)** Gene ontology analysis performed on 769 genes which are downregulated in *hyl1-2* mutant and not changed in *se-3* mutant plants.

## Discussion

The DRB1 protein plays a major role in the processing of pri-miRNAs. But it was also reported that without DRB1, in case of intron containing *MIR* genes both forms of pri-miRNA (before and after splicing) accumulate to high levels in *Arabidopsis thaliana* (Szarzynska, Sobkowiak et al. 2009). Therefore, it seems that DRB1 may have an additional function in microRNA biogenesis. It appears that the recruitment of DRB1 to the miRNA biogenesis machinery takes place at the very early stages of pri-miRNA processing, possibly before splicing occurs. For protein-coding genes, a direct connection between gene transcription and further co- and post-transcriptional processing of nascent pre-mRNAs has been shown (Bauren and Wieslander 1994, Li, Wang et al. 2019). It is known that almost all plant *MIR* genes are transcribed by RNA Pol II (Xie, Allen et al. 2005). In addition, similar to pre-mRNAs, primary transcripts of *MIR* genes also undergo further processing, including cap structure formation, polyadenylation, splicing and m^6^A methylation. (Xie, Allen et al. 2005, Bielewicz, Kalak et al. 2013, Schwab, Speth et al. 2013, Knop, Stepien et al. 2017, Bhat, Bielewicz et al. 2019). The results presented in this manuscript show that a GUS reporter under the control of a *MIR* gene promoter is expressed at a lower level when DRB1 is missing (*hyl1-2* and *hyl1-2/dcl1-13* background). Moreover, evaluation of the occupancy of RNA Pol II on *MIR* genes in *hyl1-2* showed an accumulation of total RNA Pol II in the region approximately 200 base pair upstream of the transcription initiation site. This phenomenon was observed in the cases of *MIR393A*, *MIR393B*, as well as *MIR159A, MIR163, MIR166A, MIR170* and *MIR775A* in the *hyl1-2* mutant. While the distribution of RNA Pol II in a variety of complex genomes is correlated with gene expression, the presence of RNA Pol II at a specific locus does not necessarily indicate active expression from this locus. The higher occupancy of the RNA polymerase II at the analyzed *MIR* promoter regions, together with results from the GUS reporter assays, suggest that transition of transcription from initiation to elongation is impeded in the *hyl1-2* mutant. This suggestion is supported by the results showing that DRB1 colocalized with total RNA pol II and also specifically and individually with RNA Pol II phosphorylated at Ser5 and Ser2. It is known that the phosphorylation status of the CTD domain of RNA Pol II is very important in transcription (Komarnitsky, Cho et al. 2000) and the interplay between kinases and phosphatases acting on RNA Pol II can modify gene expression. One of the proteins that is able to dephosphorylate the CTD of the RNA Pol II at Ser5 residue is a protein phosphatase called CPL1 (Koiwa, Hausmann et al. 2004). Similarly, it was reported that the phosphorylation status of DRB1 is also important in microRNA biogenesis (Mendoza, Du et al. 2005, Manavella, Hagmann et al. 2012, Raghuram, Sheikh et al. 2015, Su, Li et al. 2017, Yan, Wang et al. 2017, Achkar, Cho et al. 2018). It was shown that the DRB1 protein needs to be dephosphorylated for its optimal activity and this is also maintained by the CPL1 protein. Fully phosphorylated DRB1 exclusively localized in the nucleus, is not active in pri-miRNA processing, however this does not exclude it role in the transcription of *MIR* genes. More importantly, phosphorylation status of DRB1 could be a factor that distinguishes its role from acting in transcription and processing of pri-miRNAs. An alternative model suggests that the C-terminal region of DRB1 which displays tendencies to bind dsDNA may interact with chromatin (Bhagat, Verma et al. 2018). Our data presents new possibilities regarding DRB1’s role in the plant cell but to put forward a detailed mechanism of how DRB1 affects transcription more work.

## Material and methods

### Plant material and growth conditions

*Arabidopsis thaliana* ecotype Columbia-0 plants were used as wild-type plants, insertion mutant SALK_064863 as knockout of DRB1 *(hyl1-2)* and insertion mutant SAIL_44_G12 as *se-2. pMIR393A:GUS* and *pMIR393B:GUS* reporter lines which were used in this study were described previously (Parry, Calderon-Villalobos et al. 2009). *hyl1-2*/*dcl1-13* and p*DRB1*:*DRB1*:*HA* transgenic lines were obtained by floral dip transformation of *hyl1-2* (Clough and Bent 1998). Genomic sequences of *DCL1* and *DRB1* with around 2kb promoter region were cloned into pENTR-D-TOPO vector (Invitrogen). Next, site directed mutagenesis was performed to introduce *DCL1-13* point mutation. Finally, LR reaction was used to subcloned *DCL1-13* and *DRB1* sequence into pEarlyGate301 plasmid followed by Agrobacterium transformation (Earley, Haag et al. 2006).

Arabidopsis plants were grown in soil (Jiffy-7 42 mm; Jiffy Products International AS, Stange, Norway) or on ½-strength MS media with 0.8% agar square plates in growth chambers (Sanyo/Panasonic, Japan) that had a 16-h day length (150–200 µE/m^2^s), a constant temperature of 22 °C and 70% humidity. Seeds were sterilized before sowing in 10% sodium hypochlorite in 70% EtOH solution.

### GUS staining for reporter lines analysis

14 days old Arabidopsis seedlings which were grown in ½ MS medium were incubated in staining solution containing 1 mM X-Gluc in 100 mM Na3PO4 (pH 7.2), 0.1% Triton X-100, 5 mM K3Fe(CN)6 and 5 mM K4Fe(CN)6 for 24 h at 37°C. Seedlings were then cleared in 70% ethanol for 2 days and mounted in 50% v/v glycerol before observations. After GUS staining the pictures were taken using the Leica M60 stereo microscope.

### RNA isolation, cDNA synthesis and qPCR

Total RNA from three-week-old or 14-day-old plants was isolated using the TRIzol™ reagent (Invitrogen) and a Direct-zol RNA MiniPrep Kit (Zymo Research). The RNA was then cleaned with Turbo™ DNase (Invitrogen) according to the provided protocol. Reverse transcription reaction was performed with Superscript™ III Reverse Transcriptase (Invitrogen) and oligo-dT primer. qPCR was performed with Power SYBR™ Green PCR Master Mix (Applied Biosystems) using a QuantStudio™ 7 Flex Real-Time PCR System (Applied Biosystems). The expression levels were calculated with the 2^−ΔΔCt^ method. The Mann-Whitney U test was used for statistical analyses.

### Chromatin Immunoprecpitation

Chromatin immunoprecipitation was performed using nuclei isolated from crosslinked (1% formaldehyde) 21-day old leaves, as described in (Bowler, Benvenuto et al. 2004) with minor modifications. Sonic and IP buffer were prepared as described in (Kaufmann, Muino et al. 2010). Chromatin was sonicated at 4°C with a Diagenode Bioruptor Plus at high intensity for 30 min (30 s on/30 s off) to obtain 200-300 bp DNA fragment size. Antibodies against total RNA Pol II (Abcam ab817) were used with Dynabeads Protein G (Thermo Scientific). Protein– DNA complexes were eluted from the beads as described in (Rowley, Bohmdorfer et al. 2013) and DNA was purified using a column-based method. DNA libraries were obtained using MicroPlex Library Preparation Kit (Diagenode) and sequenced on HiSeq HO 125 SE (Fasteris). Reads were trimmed to 50bp and aligned to the Arabidopsis genome (TAIR10) using Bowtie (wit parameters -M1 –n2) (Langmead, Trapnell et al. 2009). Duplicate reads were removed by SAMtools (Li, Handsaker et al. 2009). Remaining sequences were extended to 200bp according to ChIP fragment length. Plots of RNA Pol II distribution were made using ngs.plot software (Shen, Shao et al. 2014). Genomic coordinates of *MIR* genes were taken from mirEX^2^ data base (Zielezinski, Dolata et al. 2015).

### *In situ* immunolocalization of DRB1 and RNA polymerase II

The double immunodetection experiments were performed according to the protocol described in (Bhat, Bielewicz et al. 2019). For HYL1 localization primary rabbit antibodies (Agrisera, AS06 136) diluted to 1:200 were used. After the double labeling assay, the slides were stained for DNA detection with Hoechst 33342 (Life Technology, USA) and mounted in ProLong Gold antifade reagent (Life Technologies).

### Co-immunpreciptation of partner proteins of DRB1

For each co-immunoprecipitation (co-IP), nuclear extract was prepared like described above. Proteins were extracted from nuclear pellet by resuspending it in nuclear lysis buffer (10% sucrose, 100mM Tris–HCl, pH 7.5, 5mM EDTA, 5mM EGTA, 300mM NaCl, 0.75% Triton X-100, 0.15% sodium dodecyl sulphate (SDS), 1mM dithiothreitol (DTT), and 1x cOmplete™ EDTA-free protease inhibitor (Roche)) and sonicated 2 cycle x 30 second ON/ 30 second OFF using Bioruptor® Plus (Diagenode). After removal of cell debris by centrifugation (5 min, 16000g, 4 °C) the cleared supernatants were diluted 1 time using water containing 1x cOmplete ™ EDTA-free protease inhibitor (Roche). Protein extract was incubated overnight with anti-HA antibodies (Roche). After over-night incubation, Dynabeads with protein G were added and samples were incubated for 1h. After incubation, beads were washed 2 times with low salt buffer (20mM Tris–HCl pH 8, 2mM EDTA, 150mM NaCl, 1% Triton X-100, 0.1% SDS), 1 time with high salt buffer (20mM Tris–HCl pH 8, 2mM EDTA, 500mM NaCl, 1% Triton X-100, 0.1% SDS), 1 time LiCl buffer (10mM Tris–HCl pH 8, 1mM EDTA, 250mM LiCl, 1% NP40, 0.1% SDS, 1% sodium deoxycholate) and twice with TE (100mM Tris–HCl pH 8, 1mM EDTA). Beads containing proteins were analyzed by mass spectrometry. Control IPs were performed in Col-0 using anti-HA antibodies.

Mass spectrometry analyses were performed by IBB PAS, Warsaw. Magnetic beads containing proteins were suspended in 100 mM ammonium bicarbonate buffer, reduced using 100 mM DTT for 30 min at 57 °C and alkylated in 50 mM iodoacetamide for 45 min at RT in the dark. In the next step proteins were digested overnight using 100 ng/µl trypsin (Promega) at 37°C. Peptide mixtures were separated using Nano-Ultra Performance Liquid Chromatograpy coupled to Orbitrap Velos mass spectrometer (Thermo). Peptides were identified with Mascot algorithm (Matrix Science, London, UK) and searched against the TAIR10 database. The total number of MS/MS fragmentation spectra was used to quantify each protein from at least three independent biological replicates. Biological replicates consisted of plants of the same genotype grown at different dates and in different growth chambers. For the statistical analysis we compared the data from three independent experiments for pDRB1:DRB1:HA. Statistical analysis was perform with DESEq2 R-package (Love, Huber et al. 2014).

### Analysis of RNA-seq data

RNA-seq data from wild type, *hyl1-2* and *se-3* was downloaded from published dataset under the accession number ERP001616 (Manavella, Hagmann et al. 2012). Raw reads were trimmed (first 20 nucleotides) with FASTX-Toolkit, adapters were removed using Trimmomatic and rRNA sequences where removed with Bowtie (Langmead, Trapnell et al. 2009, Bolger, Lohse et al. 2014). Clean reads were aligned to the Arabidopis TAIR10 reference genome using HISAT2 (Kim, Langmead et al. 2015). The overall alignment rate was 98-99% for each sample. Next, prepDE.py script was used to extract count information from StringTie output and DESeq2 R package was used to find differentially expressed genes (Love, Huber et al. 2014). Gene ontology was performed with clusterProfiler R package (Yu, Wang et al. 2012).

## Acknowledgments

This work was funded by the National Center of Sciences (NCN); Preludium grant UMO-2011/03/N/NZ2/03147; Sonata grant UMO-2016/23/D/NZ1/00152 and the KNOW RNA Research Centre in Poznan (01KNOW2/2014).

---

**Figure S1.**
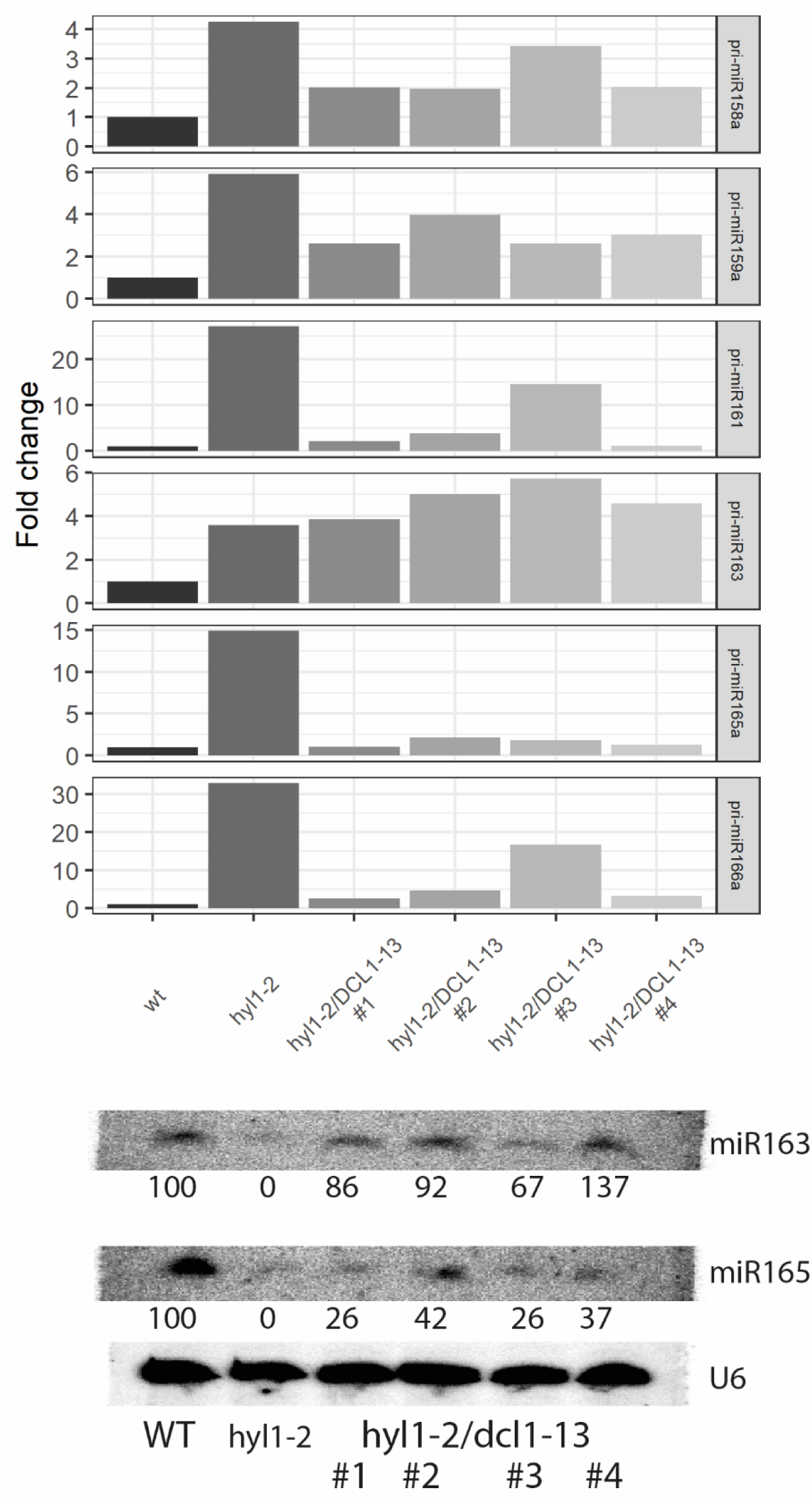
HYL1 stimulates transcription from *MIR* genes. Analysis of *hyl1-2*/*dcl1-13* transgenic lines. RT-qPCR analysis of levels of 6 pri-miRNAs in wild type, *hyl1-2* and four independent *hyl1-2*/*dcl1-13* transgenic lines (left). Error bars indicate SD (n=3). Northern blot analysis of mature microRNA levels in wild type, *hyl1-2* and four independent *hyl1-2/dcl1-13* transgenic lines (right). U6 serves as a positive control for northern blot hybridization

**Figure S2.**
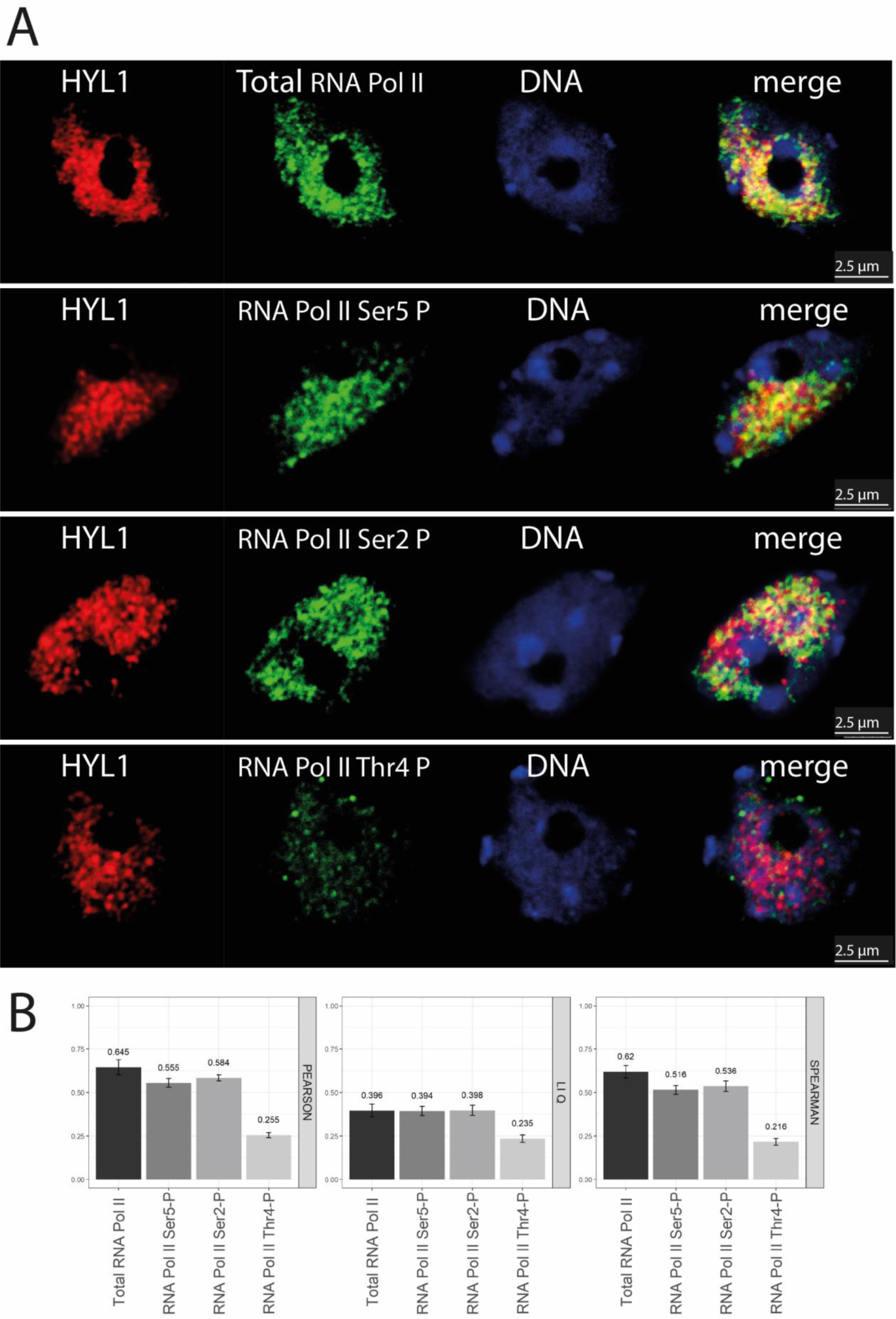
DRB1 colocalizes with RNA polymerase II in *se-2* mutant plants: **a)** Nuclei from fixed cells where DRB1 is shown in red, RNA Pol II in green and DNA in blue. The merge column shows all 3 channels. **b)** Colocalization scores of DRB1 and RNA Pol II calculated by three different approaches (Pearson, Spearman and LiQ).

**Figure S3.**
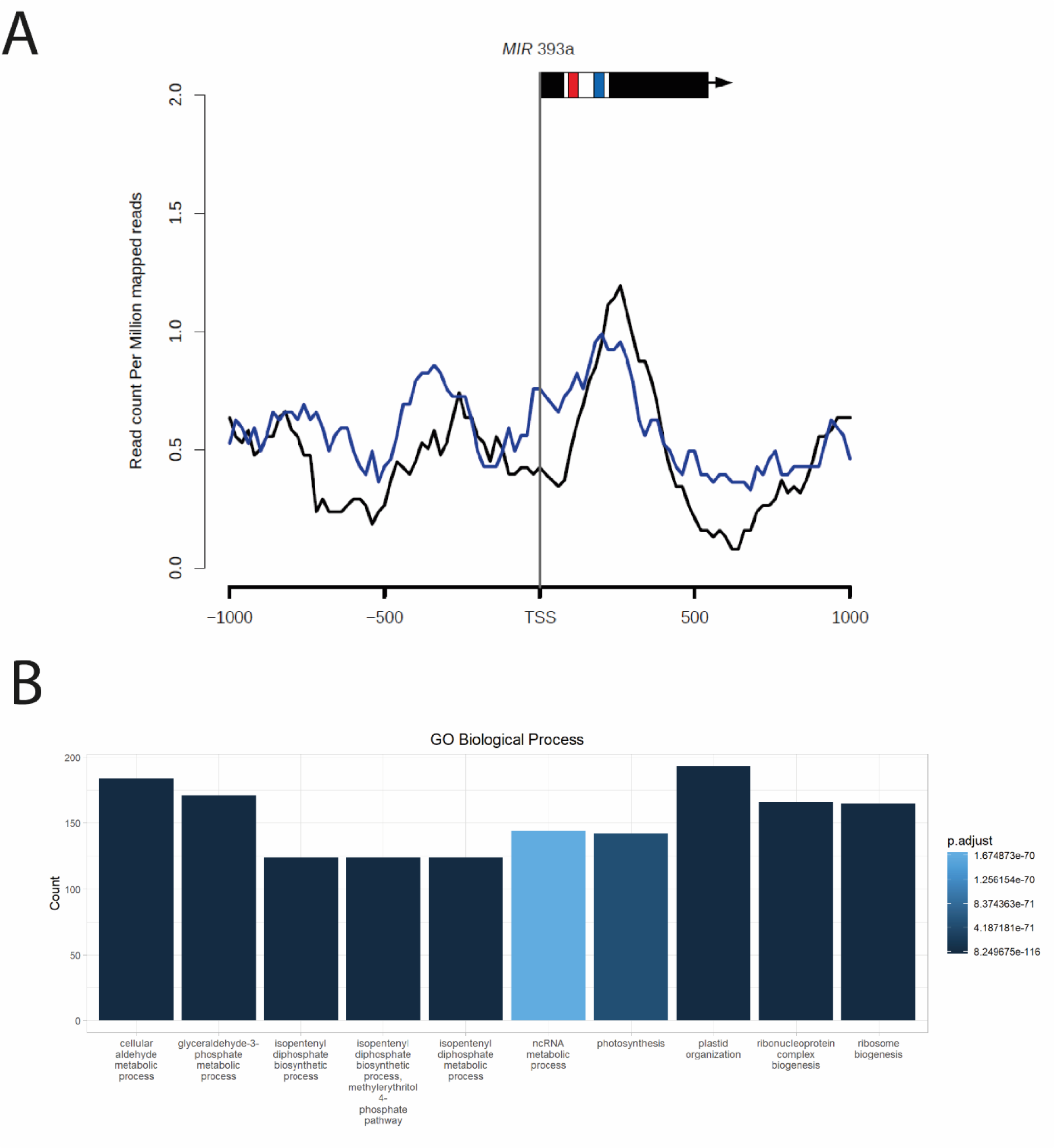
Gene ontology analysis of genes downregulated in *hyl1-2* mutant in comparison to wild type plants.

